# Investigating the causal effect of maternal vitamin B12 and folate levels on offspring birthweight

**DOI:** 10.1101/2020.01.21.914655

**Authors:** Gunn-Helen Moen, Robin N Beaumont, Christine Sommer, Beverley M. Shields, Deborah A Lawlor, Rachel M Freathy, David M Evans, Nicole M Warrington

**Affiliations:** Institute for Clinical medicine, Faculty of Medicine, University of Oslo, Norway; The University of Queensland Diamantina Institute, The University of Queensland, Woolloongabba, QLD 4102, Australia; K.G. Jebsen Center for Genetic Epidemiology, Department of Public Health and Nursing, NTNU, Norwegian University of Science and Technology, Norway; Population Health Science, Bristol Medical School, University of Bristol, UK; Institute of Biomedical and Clinical Science, College of Medicine and Health, University of Exeter, Royal Devon and Exeter Hospital, Exeter, EX2 5DW, UK; Department of Endocrinology, Morbid Obesity and Preventive Medicine, Oslo University Hospital, Norway; Medical Research Council Integrative Epidemiology Unit at the University of Bristol, Bristol, BS8 2BN, UK; Bristol National Institute of Health Research Biomedical Research Centre, UK

**Keywords:** Maternal genetic effect, Fetal genetic effect, Vitamin B12, Folate, Birthweight, Mendelian randomization

## Abstract

Lower maternal serum vitamin B12 (B12) and folate levels have been associated with lower offspring birthweight in observational studies. The aim of this study was to investigate whether this relationship is causal.

We performed two-sample Mendelian Randomization (MR) using summary data on associations between genotype-B12 (10 genetic variants) or genotype-folate (4 genetic variants) levels from a genome-wide association study of 45,576 individuals (sample 1) and maternal-specific genetic effects on offspring birthweight from the latest EGG consortium meta-analysis with 297,356 individuals reporting their own birthweight and 210,248 women reporting their offspring’s birthweight (sample 2). To investigate the effect of offspring’s own B12 or folate levels on their own birthweight, we performed two-sample MR using the fetal-specific genetic effects from the latest EGG consortium meta-analysis. We used the inverse variance weighted method, and sensitivity analyses to account for pleiotropy, in addition to sensitivity analyses excluding a potentially pleiotropic variant in the *FUT2* gene for B12.

We did not find evidence for a causal effect of maternal B12 on offspring birthweight, nor evidence for an effect of offspring B12 on their own birthweight using the fetal-specific genetic effect. The results were consistent across the different methods and in sensitivity analyses excluding the *FUT2* variant. We found a positive effect of maternal folate on offspring birthweight (0.146 [0.065, 0.227], which corresponds to an increase in birthweight of 71g per 1SD higher folate). We found some evidence for a small inverse effect of fetal folate on their own birthweight (−0.051 [−0.100, −0.003]).

In conclusion, our results are consistent with evidence from randomized controlled trials that increased maternal folate levels increase offspring birthweight. We did not find evidence for a causal effect of B12 on offspring birthweight, suggesting previous observational studies may have been due to confounding.

## Introduction

Low birthweight is not only associated with lower infant survival, but also increased future risk of chronic diseases in offspring such as type two diabetes mellitus and cardiovascular disease. The Developmental Origins of Health and Disease (DOHaD) hypothesis provides an explanation for these robust observational associations[1–7], which stipulates that impaired fetal growth and development *in utero* leads to developmental compensations that program the offspring to increased risk of disease in later life[5]. In other words, maternal under- or over-nutrition during pregnancy not only exerts detrimental effects on offspring birthweight, but is hypothesized to cause long term effects on the cardio-metabolic health of the offspring. Therefore, identifying causal relationships between maternal nutrition and offspring birthweight could provide information on modifiable maternal exposures to ensure that more babies are born within a healthy weight range.

Low maternal folate (vitamin B9) and vitamin B12 levels have previously been linked with low offspring birthweight and increased risk of preterm birth [8–12]. The combination of high folate and low vitamin B12 levels have also been associated with low birthweight[13]. However, the association between B12 and offspring birthweight has been questioned, with a systematic review and meta-analysis showing no evidence for an association (5.1g change in birthweight per 1 standard deviation (SD) increase in B12 [95% confidence interval: −10.9g, 21.0g]) [14]. For folate, data from a meta-analysis suggested a 2% (95% confidence interval: 0.7%, 3.5%) higher birthweight for every two-fold higher folate intake[9] and a Cochrane review suggested a 135g (95% confidence interval: 47.85g, 223.68g) increase in birthweight with folic acid supplement use[15]. However, the observational association between maternal folate and B12 and offspring birthweight could be confounded by other maternal characteristics, such as high body mass index (BMI) and gestational diabetes.

To ensure pregnant women have adequate levels of folate during their pregnancy to prevent neural tube defects, folic acid supplements are recommended to all women planning pregnancy and during the antenatal period. In addition, many countries fortify certain foods with folic acid. For vitamin B12 recommendations vary, with most high income countries not recommending supplements unless there is clear deficiency. Folate and B12, along with vitamin B6 and B2, are coenzymes, or integral components of coenzymes, that are involved in one carbon metabolism, also known as the folate metabolism pathway [16]. One carbon metabolism is essential for DNA synthesis and DNA methylation [16, 17], which is particularly important during pregnancy because of the cell division and differentiation occurring *in utero*. The interplay between folate and B12 is important both because a folate deficiency can be induced by a B12 deficiency [18], and because high folate levels can mask a B12 deficiency [19]. The interrelationship between these two vitamins is best explained by the methyl trap hypothesis. This hypothesis states that a vitamin B12 deficiency can lead to lowered levels of methionine synthetase, which results in a functional folate deficiency by trapping an increased proportion of folate as the 5-methyl derivative [18]. Low folate and B12 levels may be caused by low dietary intake, lack of intrinsic factor needed to absorb vitamin B12, poor absorption of ingested folate/B12, or an alteration of folate metabolism due to genetic defects or drug interactions.

Conclusions regarding causality cannot easily be drawn from observational multivariable regression association studies. Mendelian randomization (MR) is a method that uses genetic variants as instrumental variables to provide information on causality in observational studies [20]. Offspring birthweight is thought to be influenced by offspring genetics, maternal genetics operating on the intrauterine environment, and environmental factors. MR investigating the effect of maternal environmental exposures on offspring birthweight is therefore more complicated because we have both maternal and offspring genetics (which are correlated, r≈0.5) influencing the outcome of interest. A recently described statistical method based on structural equation modelling can be used to estimate the independent effect of maternal genotypes on offspring outcomes by conditioning on offspring genotype[21]. These maternal-specific genetic effects can then be used in a two-sample MR framework to estimate the causal effect of maternal environmental exposures, such as B12 or folate, on offspring birthweight [22]. The aim of this study was to investigate whether there is a causal relationship between serum levels of maternal B12 or folate and offspring birthweight.

## Methods

### Exposure measurements

We extracted summary results data on 10 single nucleotide polymorphisms (SNPs) that were robustly associated with serum B12 and four SNPs that were robustly associated with folate levels in the largest genome-wide association study (GWAS) of these phenotypes to date[23] (n=45,576 individuals of Danish and Icelandic ancestry; Supplementary Table 1). We only had access to the SNPs that were deemed significant (P<1×10^−8^ in the combined Icelandic and Danish analysis), so we were unable to look at the effect of the folate SNPs on B12 and vice versa, or undertake multivariable Mendelian Randomization to explore independent (of each other) effects of folate and B12 on birthweight. The *FUT6* variant, rs7788053, was identified in the GWAS as associated with B12 but was not included in our analysis as it is monoallelic in the CEU (Utah Residents from North and West Europe (https://ldlink.nci.nih.gov/)) population. The SNP was not available in the GWAS of birthweight and therefore left out of the analysis.

### Outcome measurements

We obtained summary results statistics from the latest GWAS of birthweight published by the Early Growth Genetics (EGG) consortium[24], which included full term births (defined as 37 or more weeks of gestation where gestational age was available, or a birthweight between 2.5-4.5 kg for UK Biobank). The GWAS of birthweight included 297,356 individuals reporting their own birthweight and 210,248 women reporting their offspring’s birthweight. Due to the correlation between maternal and fetal genotypes, and the influence of both genotypes on offspring birthweight, the authors used a structural equation model[21] to estimate the independent maternal and fetal genetic effects on birthweight (this model is similar to a conditional linear model accounting for both maternal and fetal genotype). We extracted the summary results statistics for our B12 and folate SNPs from this GWAS and used the maternal- and fetal-specific effect estimates in the MR analysis (Supplementary Table 2). The *MMACHC* SNP, rs12272669, which is associated with B12 was not available in the GWAS summary results statistics for birthweight, so we extracted a SNP in high LD, rs11234541 (R^2^=0.942, D’=1 (https://ldlink.nci.nih.gov/)).

In the folate GWAS, the *MTHFR* SNP, rs17421511, was identified to be associated with folate in an analysis that conditioned on another *MTHFR* SNP, rs1801133. Therefore, we performed an approximate conditional and joint (COJO) analysis [25] in GCTA[26] using European ancestry meta-analysis summary statistics for the maternal- and fetal-specific effects on birthweight, conditioning on rs1801133. The reference panel for determining linkage disequilibrium (LD) was made up of 47,674 unrelated UK Biobank participants defined by the UK Biobank as having British ancestry [27]. We extracted the maternal- and fetal-specific effect estimates for rs17421511 from the COJO analysis (Supplementary Table 2).

### Statistical analysis

To assess instrument strength for the standard inverse variance weighted (IVW) MR analysis, we calculated the approximate F statistics for each of the folate and B12 SNPs that will be used in our two-sample MR study (i.e., F ≈ (β/SE)^2^) (Supplementary table 1). In the case of MR Egger regression analysis [28], the F statistics from the individual markers are not a sufficient indicator of instrument strength. Rather Bowden and colleagues [29] show that an I^2^_GX_ statistic can be used to quantify instrument strength in MR Egger analyses. The authors show that a high value of I^2^_GX_ (i.e. close to one) suggests that the instrument effect sizes are estimated well and that measurement error/weak instrument bias is unlikely to affect the results of standard MR Egger analyses. On the other hand, if I^2^_GX_ is less than 0.9, inference from MR Egger should be interpreted with caution and some alternative sensitivity analyses should be considered. As such, we calculated I^2^_GX_ values for our SNPs to assess the instrument strength in the MR Egger analysis.

We performed two-sample, IVW MR analysis to estimate the causal effect of maternal B12 or folate levels on offspring birthweight. We performed a secondary analysis investigating the causal relationship between an individual’s own B12 or folate levels and their own birthweight using the results of the fetal-specific genetic effects on birthweight. Analyses were performed with the TwoSampleMR package[30] (https://github.com/MRCIEU/TwoSampleMR) in R version 3.5.2 (https://cran.r-project.org/). We performed a test of heterogeneity of causal effect estimates across each of the SNPs using Cochran’s Q. If heterogeneity was detected, leave-one-out IVW analysis was performed to assess the effect of the individual SNPs on the overall causal estimate. We tested for directional pleiotropy using the MR Egger intercept[28]. Additionally, we conducted sensitivity analysis to adjust for any directional pleiotropy using MR Egger regression[28], weighted median [31], simple and weighted mode estimation [32]. We acknowledge that these sensitivity analyses perform well with large numbers of SNPs and may not perform well in our analyses for effects of folate where we only have four SNPs; however, we include the results of the MR Egger for completeness. As the weighted and simple mode estimator select the group of SNPs that produce the most common causal effect on the outcome we chose not to perform these analysis for folate where only four SNPs were available.

Further sensitivity analyses were conducted for the B12 analysis excluding the *FUT2* variant, rs602662. The FUT2 protein, which is expressed in the small intestine, is a part of the protein glycosylation pathway and is involved in the absorption and modification of several nutrients [33–35]. Therefore, it may reflect nutritional status, which in turn could affect a multitude of lifestyle-associated traits and diseases. In a previous MR study investigating the causal relationship between B12 and BMI[36]; the causal relationship identified between low B12 levels and higher BMI disappeared when the variant in the *FUT2* gene was excluded from the analysis [36]. We therefore performed the MR analyses both with and without the *FUT2* variant to assess the effect of this potentially pleiotropic variant on the causal estimate, as has been done in previous papers[36, 37].

Three of the variants associated with folate levels were located within the *MTHFR* gene. We looked up the LD between the variants (Supplementary table 3) and conducted a leave-one-out MR analysis as a sensitivity analysis to assess the influence of the individual SNPs.

### Follow-up analysis of gestational duration

Gestational duration is a major determinant of birthweight. Since gestational duration was unavailable for >85% of individuals in the GWAS of birthweight[24], it is possible that a causal relationship between maternal B12 or folate and birthweight could be driven by a relationship with gestational duration. Therefore, we performed a two-sample, IVW MR analysis to estimate the causal effect of maternal B12 and folate levels on gestational duration. We obtained summary results statistics from the latest maternal GWAS study of gestational duration of 43,568 women of European ancestry from Zhang *et al*[38]. For the fetal GWAS of gestational duration we obtained summary results statistics from the latest EGG Consortium study of 84,689 individuals [39]. For the B12 analysis the *MMACHC* SNP rs12272669 was not available in the maternal or fetal gestational duration GWAS (and no SNPs in high LD (R^2^ > 0.8) were available), leaving a total of nine SNPs for analysis. The effect allele frequencies were not available in the publicly available summary statistics from the fetal GWAS of gestational duration, so we removed the *MMAA* SNP, rs2270665, from the B12 analysis as it is a palindromic SNP and we could not confirm that the direction of effect was the same as in the B12 summary statistics. We conducted sensitivity analysis using MR Egger regression[28], weighted median [31], simple and weighted mode estimation [32] to adjust for any potentially pleiotropic SNPs. Similar to the birthweight analysis, we caution the interpretation of the folate sensitivity analyses due to the small number of SNPs available.

### Checking SNP associations with B12 and folate levels in pregnant women from the EFSOCH study

We performed an association analysis of the B12/folate SNPs and the serum level of the two vitamins, measured at 28 weeks of gestation [40] in a total of 871 pregnant women from The Exeter Family Study of Childhood Health (EFSOCH)[41]. Genome-wide genotyping and imputation in this study have been described previously[42]. The outcome variables were inverse-normal transformed before analysis, and the linear regression analysis was adjusted for maternal age and five ancestry principal components. The rs12272669 variant in the *MMACHC* gene was not available, so rs11234541 in high LD (r^2^=0.9421 in CEU) was used as a proxy. This SNP had low imputation quality in the EFSOCH study (INFO=0.68), and the results should be interpreted with care.

### Informed consent

Only publically available summary results statistics were used in the birthweight MR analyses. Participants in EFSOCH and 23andMe provided informed consent to participate in those respective cohort. Summary data was provided to us for further analysis.

## Results

The 10 variants included in the MR analysis of maternal serum B12 explained 5.24% of the variance in serum B12 concentrations in the original GWAS (3.82% without the *FUT2* variant, rs602662). The four variants included in the MR analysis of maternal folate explained 1.3% of the total variation in serum folate concentration in the original GWAS. F statistics for individual SNPs ranged from 35 to 623 for B12 and 48 to 203 for folate. The magnitude of the SNP effects on serum B12 and folate concentrations measured in pregnant women from the EFSOCH study were consistent with those from the original GWAS (Supplementary Figure 3 and 4), apart from the *MMACHC* variant, rs12272669, which appeared to have little effect in pregnant women at 28 weeks gestation (heterogeneity P=0.0002). However, we used a proxy for this variant in the EFSOCH study, rs11234541, which was in high LD with rs12272669 but had low imputation quality.

The causal effect estimates of maternal and fetal B12 and folate on birthweight from the IVW analysis and sensitivity analyses are shown in Table 1. We found no evidence for a causal effect of maternal B12 levels on offspring birthweight (0.009 SD change in birthweight per 1SD higher B12 (SE_IVW_=0.012), P_IVW_=0.469; Figure 1), or of fetal B12 levels on their own birthweight (−0.012 SD change in birthweight per 1SD higher B12 (SE_IVW_=0.017), P_IVW_=0.478; Supplementary Figure 1). The effect estimates were similar when excluding the *FUT2* variant, rs602662 (Supplementary Table 4).

**Table 1:**
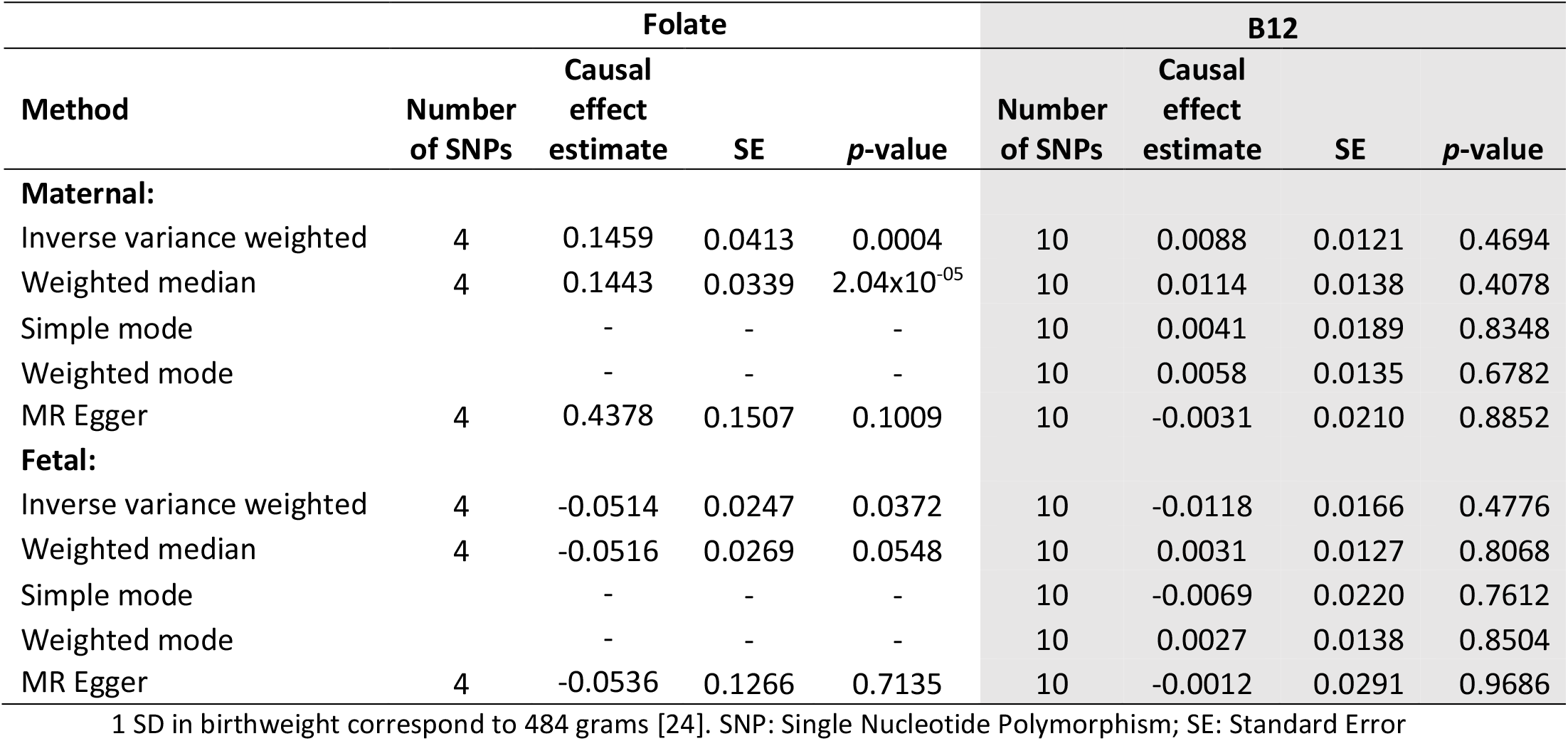
Mendelian randomization estimates of the causal effect of folate and vitamin B12 levels on birthweight. We estimated causal effects for maternal folate and B12 levels on offspring birthweight (Maternal) and offspring’s folate and B12 levels on their own birthweight (Fetal). Causal effects are estimated using five MR models: inverse variance weighted (IVW), weighted median, simple mode, weighted mode and MR Egger regression. Causal effect estimates are the difference in mean birthweight (in standard deviation; SD) per 1SD higher folate/B12.

| Method | Folate |  |  |  | B12 |  |  |  |
| --- | --- | --- | --- | --- | --- | --- | --- | --- |
|  | Number of SNPs | Causal effect estimate | SE | p-value | Number of SNPs | Causal effect estimate | SE | p-value |
| **Maternal:** |  |  |  |  |  |  |  |  |
| Inverse variance weighted | 4 | 0.1459 | 0.0413 | 0.0004 | 10 | 0.0088 | 0.0121 | 0.4694 |
| Weighted median | 4 | 0.1443 | 0.0339 | $2.04 \times 10^{-05}$ | 10 | 0.0114 | 0.0138 | 0.4078 |
| Simple mode |  | - | - | - | 10 | 0.0041 | 0.0189 | 0.8348 |
| Weighted mode |  | - | - | - | 10 | 0.0058 | 0.0135 | 0.6782 |
| MR Egger | 4 | 0.4378 | 0.1507 | 0.1009 | 10 | -0.0031 | 0.0210 | 0.8852 |
| **Fetal:** |  |  |  |  |  |  |  |  |
| Inverse variance weighted | 4 | -0.0514 | 0.0247 | 0.0372 | 10 | -0.0118 | 0.0166 | 0.4776 |
| Weighted median | 4 | -0.0516 | 0.0269 | 0.0548 | 10 | 0.0031 | 0.0127 | 0.8068 |
| Simple mode |  | - | - | - | 10 | -0.0069 | 0.0220 | 0.7612 |
| Weighted mode |  | - | - | - | 10 | 0.0027 | 0.0138 | 0.8504 |
| MR Egger | 4 | -0.0536 | 0.1266 | 0.7135 | 10 | -0.0012 | 0.0291 | 0.9686 |

1 SD in birthweight correspond to 484 grams [24]. SNP: Single Nucleotide Polymorphism; SE: Standard Error

**Figure 1:**
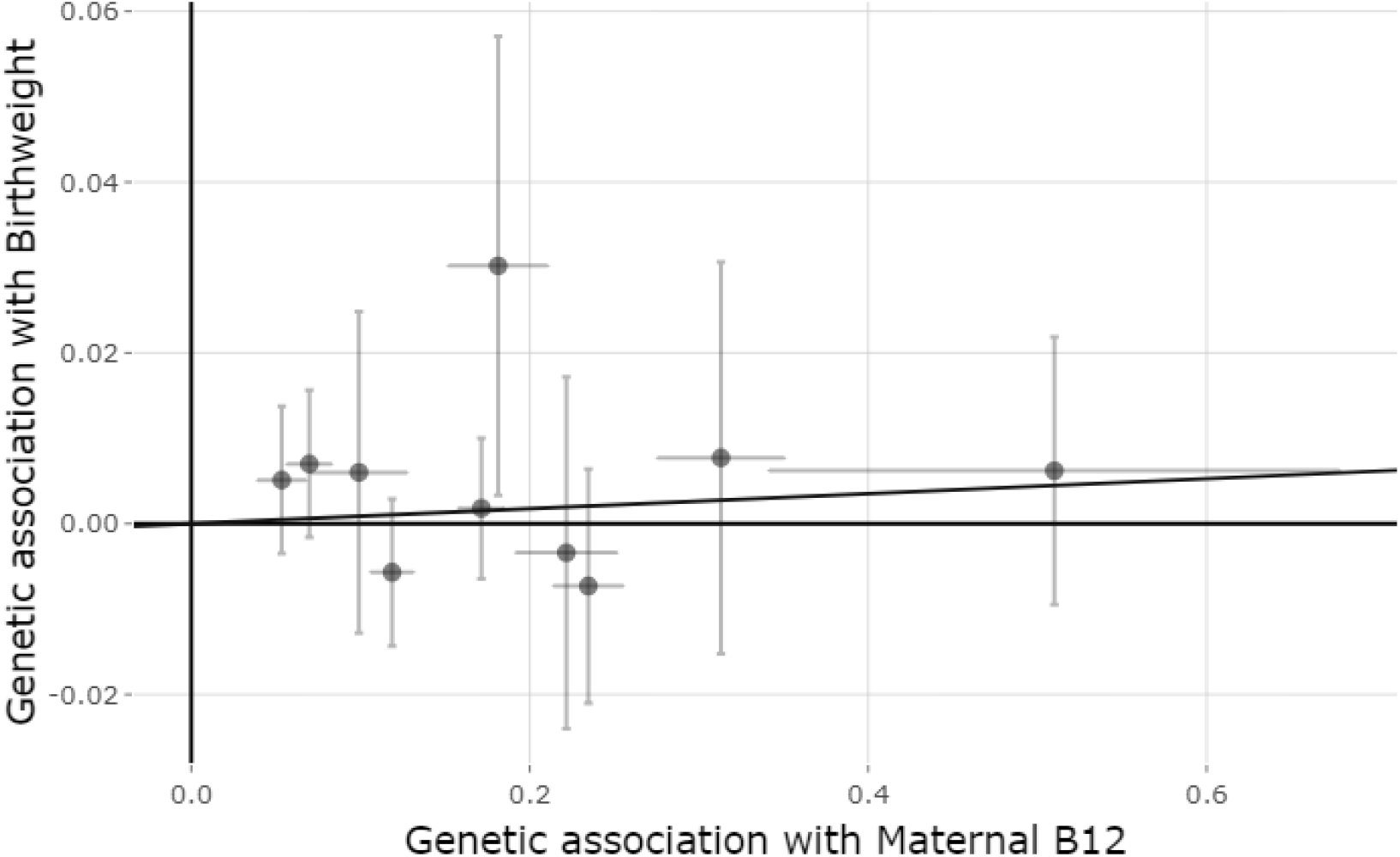
Mendelian randomization estimates of the causal effect of maternal B12 levels on offspring birthweight. X-axis shows the SNP effect, and standard error, on vitamin B12 levels for each of the 10 SNPs and Y-axis shows the SNP effect, and standard error, on offspring birthweight. The regression line for the IVW MR method is shown.

We found evidence of a positive causal effect of maternal folate levels on offspring birthweight (0.146 SD change in birthweight per 1SD higher folate (SE_IVW_=0.041), P_IVW_=4×10^−4^; Figure 2). Furthermore, we identified a negative effect of fetal folate on their own birthweight, although the effect size was nearly a third of the maternal folate causal effect estimate (−0.051 SD change in birthweight per 1SD higher folate (SE_IVW_=0.025), P_IVW_=0.037; Supplementary Figure 2).

**Figure 2:**
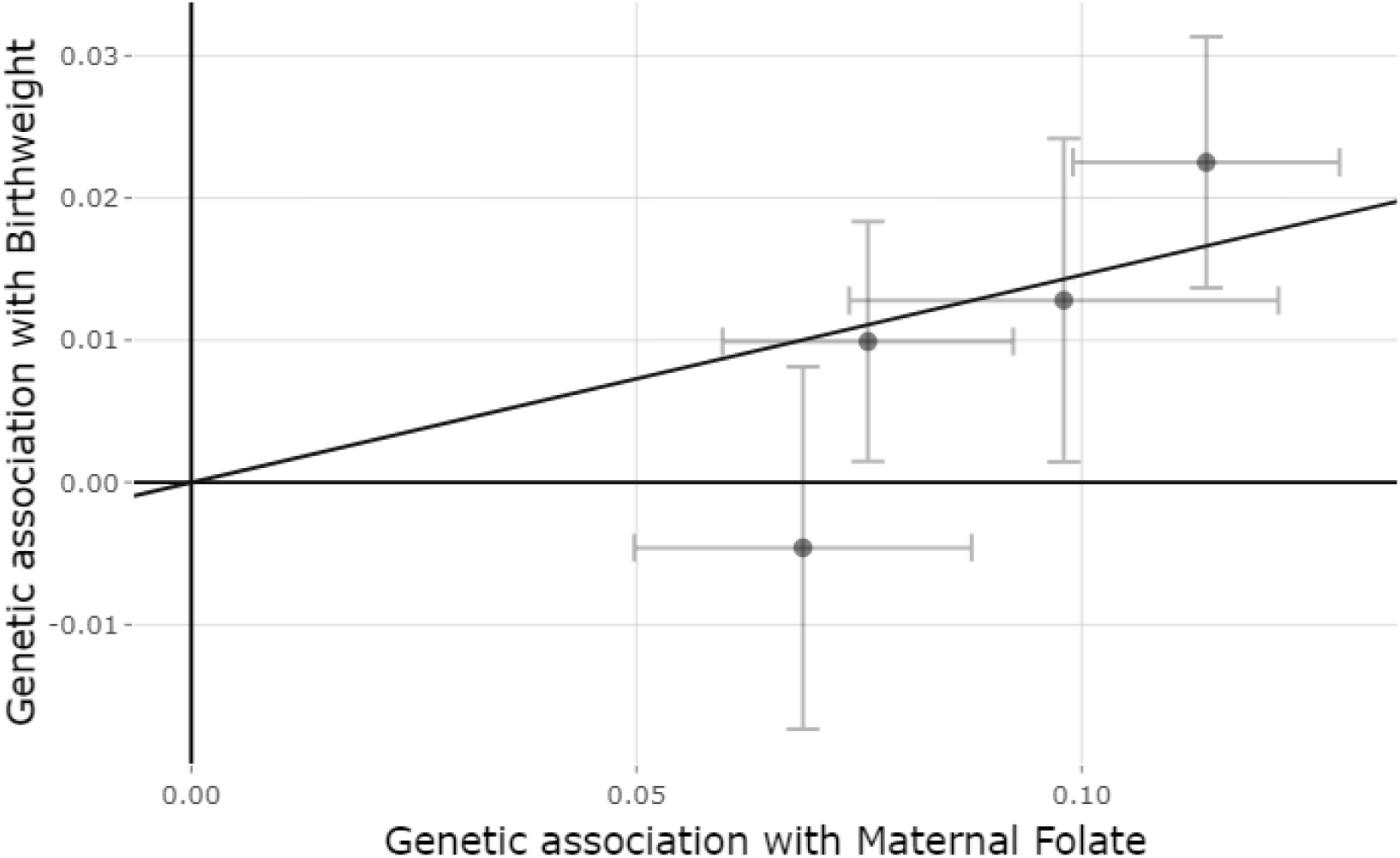
Mendelian randomization estimates of the causal effect of maternal folate levels on offspring birthweight. X-axis shows the SNP effect, and standard error, on folate levels for each of the four SNPs and Y-axis shows the SNP effect, and standard error, on offspring birthweight. The regression line for the IVW MR method is shown.

There was evidence of heterogeneity in the causal effect estimates of fetal B12 levels on their own birthweight across the individuals SNPs (Q=27.790, P=0.001), but no strong evidence of heterogeneity in the causal effect estimates of maternal B12 levels on offspring birthweight (Q=12.486, P=0.162). To investigate the heterogeneity further we performed leave one out analysis of fetal B12 levels on their own birthweight (Supplementary Table 5). As indicated by the heterogeneity test, there was some variability in the causal estimates, particularly when leaving the *MMACHC* SNP, rs12272669, or the *FUT2* SNP, rs602662, out of the analysis. Even though there are no known pleiotropic pathways for the *MMACHC* gene, we a priori expected the *FUT2* gene to have pleiotropic effects [36, 37]. The causal effect estimates of maternal folate on offspring birthweight across the individual SNPs also showed some heterogeneity (Q=6.904, P=0.075), but no evidence for the fetal folate levels (Q=0.018, P=0.999). To investigate the potential influence of the SNPs in LD in the *MTHFR* gene and potential heterogeneity identified for the maternal folate levels, we performed a leave one out analysis on both maternal and fetal folate levels on birthweight (Supplementary Table 6). When the *MTHFR* SNP rs1801133 was excluded from the analysis the causal effect was attenuated for the effect of maternal folate levels (0.099, SE=0.051, P=0.051), but stayed the same and for fetal folate levels (−0.051, SE=0.034, P=0.132).

There was no evidence of directional pleiotropy from the MR Egger regression analysis for either B12 or folate levels on birthweight (Egger intercept for maternal B12 levels = 0.003, P=0.500; Egger intercept for fetal B12 levels = −0.002, P=0.662; Egger intercept for maternal folate levels = −0.028, P=0.187; Egger intercept for fetal folate levels = 0.000, P=0.988), although again highlighting that the MR Egger result from the folate analysis would be unreliable due to the small number of SNPs. This is further supported by the low *I*^2^_GX_ of 0.83 for the folate analysis, which is below the suggested cut-off of 0.9, suggesting that the MR Egger results could be influenced by measurement error or weak instrument bias. Bowden’s *I*^2^_GX_ was 0.98 for B12 suggesting little influence in the MR Egger analyses from measurement error or weak instrument bias

Lastly, we did not find any evidence of a causal effect of maternal or fetal B12 or folate levels on gestational duration (Table 2). There was no evidence of directional pleiotropy from the MR Egger regression analysis or heterogeneity between the causal estimates at each SNP (Maternal: Directional pleiotropy: P_B12_=0.28, P_Folate_=0.87; Heterogeneity: P_B12_=0.08, P_Folate_=0.63; Fetal: Directional pleiotropy: P_B12_=0.59, P_Folate_=0.38; Heterogeneity: P_B12_=0.12, P_Folate_=0.21).

**Table 2:** Causal effect estimates of maternal folate and vitamin B12 levels on gestational duration. Causal effects are estimated using five MR models: inverse variance weighted (IVW), weighted median, simple mode, weighted mode and MR Egger regression. Causal effect estimates are the difference in mean gestational duration (in days for the maternal effect and SD for the fetal effect) per 1SD higher folate/B12.

| Method | Folate |  |  |  | B12 |  |  |  |
| --- | --- | --- | --- | --- | --- | --- | --- | --- |
|  | Number of SNPs | Causal effect estimate* | SE | p-value | Number of SNPs | Causal effect estimate* | SE | p-value |
| **Maternal:** |  |  |  |  |  |  |  |  |
| Inverse variance weighted | 4 | -0.0772 | 0.1065 | 0.4685 | 9 | -0.0683 | 0.0725 | 0.3457 |
| Weighted median | 4 | -0.1248 | 0.1261 | 0.3225 | 9 | -0.0993 | 0.0748 | 0.1844 |
| Simple mode | - | - | - | - | 9 | -0.1760 | 0.1349 | 0.2281 |
| Weighted mode | - | - | - | - | 9 | -0.1171 | 0.0885 | 0.2222 |
| MR Egger | 4 | -0.1742 | 0.5498 | 0.7813 | 9 | -0.2231 | 0.1512 | 0.1836 |
| **Fetal:** |  |  |  |  |  |  |  |  |
| Inverse variance weighted | 4 | -0.0233 | 0.0459 | 0.6101 | 8 | 0.0132 | 0.0195 | 0.5007 |
| Weighted median | 4 | -0.0700 | 0.0475 | 0.1407 | 8 | 0.0194 | 0.0212 | 0.3595 |
| Simple mode | - | - | - | - | 8 | 0.0171 | 0.0347 | 0.6363 |
| Weighted mode | - | - | - | - | 8 | 0.0279 | 0.0270 | 0.3362 |
| MR Egger | 4 | 0.2572 | 0.2561 | 0.4209 | 8 | 0.0354 | 0.0444 | 0.4557 |

\*Note that these estimates are not maternal- and fetal-specific estimates derived from conditional analyses.
SNP: Single Nucleotide Polymorphism; SE: Standard Error

## Discussion

We did not find any evidence for a causal effect of maternal B12 levels on offspring birthweight (0.009 [−0.015, 0.033] SD change in birthweight per 1SD higher B12, corresponding to a 4g increase [−7g,16g] in birthweight per 1SD higher B12), nor was there evidence that fetal B12 levels influenced their own birthweight. The results were consistent across the different sensitivity methods applied, including the analyses with or without the *FUT2* variant. In contrast, we found evidence of a positive causal effect of maternal folate levels on offspring birthweight (0.146 [0.065, 0.227] SD change in birthweight per 1SD higher folate, which corresponds to an increase in birthweight of 71g [31g,110g] per 1SD higher folate). Although we could not detect any evidence for directional pleiotropy or heterogeneity in these analysis, we have limited capacity to test this due to having only four SNPs associated with folate. We also found some evidence of an inverse effect of fetal folate levels on their own birthweight (−0.051 [−0.100, −0.003] SD change in birthweight per 1SD higher folate, corresponding to a 25g [−48g,1g] decrease in birthweight per 1SD higher folate).

Previous multivariable regression association studies investigating the relationship between maternal B12 levels and offspring birthweight have been inconsistent[8]. Nevertheless, these studies are all relatively small (samples sizes from 51 to 5,577 in the meta-analysis presented by Rogne and colleagues [8]) and therefore the effect size estimates from the multivariable regression analyses between maternal B12 levels and offspring birthweight had large confidence intervals. In addition, the meta-analysis combining the results from 12 studies found no evidence of an association between maternal B12 levels and offspring birthweight (N=9,406), with a similar estimate to the results of our MR analysis (5.1g [−10.9g, 21.0g] change in birthweight per 1SD increase in maternal B12 levels). Multivariable regression association studies and MR have different assumptions and sources of bias; however, with regards to maternal B12 levels on offspring birthweight, the effect estimates in the meta-analysis of multivariable regression analyses and our MR are similar, providing additional confidence in our result. In the meta-analysis, Rogne and colleagues [8] observed some heterogeneity between the studies (I^2^ = 30%)[8], which was driven by an association between B12 deficiency and birthweight in low and middle income countries. They suggested that this association could be explained by preterm birth rather than reduced fetal growth; however, we did not see any evidence for this in our MR analysis with gestational duration. Notably, one of the 10 SNPs from the birthweight analysis was not available for the MR analysis estimating the causal effect of maternal B12 levels on gestational duration. Importantly, even though we could not find any evidence of an effect of B12 levels on offspring birthweight within the normal range, as folate deficiency can be induced by a B12 deficiency [18] we cannot rule out an indirect effect of B12 on birthweight through folate levels. As such, monitoring both B12 and folate levels during pregnancy might be important.

Consistent with previous multivariable regression association studies[9] and randomized controlled trials (RCTs)[15], our MR study suggests a causal effect of increased folate levels on increased birthweight. Our MR analysis goes one step further than the multivariable regression studies and RCTs as we have used serum levels of folate whereas the previous studies were based on the use of folic acid supplements in pregnancy without quantifying the downstream effect that those supplements have on serum levels[9, 15]. We found a potential negative causal effect of fetal folate levels as proxied by the fetal genotype on birthweight, although the effect size was only about a third of the positive effect of maternal folate levels on offspring birthweight. The mechanism underlying this effect is unclear as folate is actively transported across the placenta by folate transporters, and not produced by the fetus [43, 44]. However, folate is metabolised into the active forms both in mothers and in the fetus, so this negative causal effect might indicate that fetal folate metabolism is important. There were only four SNPs included in the folate analysis, three of which are located in the same gene, which was not enough to obtain reliable sensitivity analyses, as partly indicated by the *I*^2^_GX_ of 0.83 for the MR Egger regression. Furthermore, the variant rs1801133 in the *MTHFR* gene has been associated with diastolic blood pressure, which is also known to causally influence birthweight[24] and could therefore indicate potentially pleiotropic pathways (Causal effect estimate from the leave-one-out analysis removing rs1801133: 48g [0g,96g] increase in birthweight per 1SD higher folate).

We did not find any evidence of a causal effect of maternal or fetal folate levels on gestational duration, suggesting that our finding of a causal effect of folate on offspring birthweight cannot be attributed to gestational duration. However, the sample size in the gestational duration GWAS (Maternal: N=43,568; Fetal: N=84,689) was smaller than the GWAS of birthweight (N=297,356 individuals reporting their own birthweight and 210,248 women reporting their offspring’s birthweight) and the maternal SNP effect estimates on gestational duration were not conditioned on fetal genotype.

It is important to examine the findings of a causal relationship between folate and birthweight within the bigger picture. Given that low maternal folate levels during pregnancy are known to cause neural tube defects[45], we would obviously not argue for stopping the widespread recommendation and use of folic acid supplements during pregnancy and the fortification of foods. Indeed, based on our results, folic acid supplementation and fortification of foods may have contributed to the observed increase in birthweight and reduction of babies born small for gestational age seen over the past few decades (23g to 46g - in a 9-18 year time span)[46–48].

### Strengths and limitations

One of the strengths of our study was the ability to use the partitioned maternal and fetal genetic effects from the largest GWAS of birthweight (N=297,356 individuals reporting their own birthweight and 210,248 women reporting their offspring’s birthweight) in a two sample MR approach to estimate the effect of maternal B12 and folate levels on offspring birthweight, independent of any direct fetal effect. We performed a series of sensitivity analyses to explore potential bias due to horizontal pleiotropy, including comparing findings for B12 levels with and without the *FUT2* genetic instrument which has been shown to have pleiotropic effects. However, we acknowledge that the methods used for the sensitivity analyses work best with more SNPs than we had available in our analyses. For folate, the *I*^2^_GX_ of 0.83, below the suggested cut-off of 0.9 indicate that the MR Egger results could be influenced by measurement error or weak instrument bias.

An important limitation with this study is that the folate and vitamin B12 levels measured and used in the exposure GWAS was from non-pregnant women, as well as men. However, we were able to check that the instruments were valid in a pregnant population by performing an association analysis in EFSOCH study. We found no evidence of different effect sizes in pregnant women compared to the exposure GWAS used in these analyses – except for in the rs12272669 variant in the *MMACHC* gene. The proxy SNP we used in the EFSOCH study, rs11234541, had low imputation quality, and as such the results should be interpreted with care.

Lastly, it is important to note that even though we did not find any effect of maternal B12 levels on offspring birthweight that does not mean that B12 levels do not have an effect on the development of the fetus. B12 levels could influence growth in a particular period of the pregnancy, effect a particular organ, or growth of fat, lean or skeletal tissue in different directions, which may not be reflected in an outcome such as birthweight.

## Conclusion

In conclusion, evidence is lacking for a causal relationship of maternal vitamin B12 levels on offspring birthweight, indicating that the association detected in previous multivariable regression association studies may have been due to confounding. We found evidence for a causal effect of increased maternal folate levels increasing offspring birthweight, which is consistent with previous multivariable regression association studies studies and RCTs.

## Acknowledgments

This article has been accepted for publication in the International Journal of Epidemiology, published by Oxford University Press.

This research was carried out at the Translational Research Institute, Woolloongabba, QLD 4102, Australia. The Translational Research Institute is supported by a grant from the Australian Government. Support for this research have been given by the Norwegian Diabetes Association and Nils Normans minnegave. We would also like to thank the research participants and employees of 23andMe for making data on maternal gestational duration available. Data on fetal gestational duration has been contributed by the EGG Consortium and the iPSYCH Consortium and has been downloaded from www.egg-consortium.org. G.H.M is supported by the Norwegian Research Council (Post doctorial mobility research grant 287198). D.M.E. is funded by an Australian National Health and Medical Research Council Senior Research Fellowship (APP1137714) and NHMRC project grants (GNT1125200, GNT1157714). R.M.F. and R.N.B. are supported by Sir Henry Dale Fellowship (Wellcome Trust and Royal Society grant: WT104150). N.M.W. is supported by an Australian National Health and Medical Research Council Early Career Fellowship (APP1104818). D.A.L’s contribution to this work is supported by the European Research Council (DevelopObese; 669545), US National Institute for Health (R01 DK10324) and the UK National Institute of Health Research (NF-0616-10102). D.A.L and D.M.E work in or are affiliated with the Medical Research Council Integrative Epidemiology Unit, which is supported by the University of Bristol and UK Medical Research Council (MC_UU_00011/6). The Exeter Family Study of Childhood Health (EFSOCH) was supported by South West NHS Research and Development, Exeter NHS Research and Development, the Darlington Trust and the Peninsula National Institute of Health Research (NIHR) Clinical Research Facility at the University of Exeter. The opinions given in this paper do not necessarily represent those of NIHR, the NHS or the Department of Health. Genotyping of the EFSOCH study samples was funded by the Welcome Trust and Royal Society grant 104150/Z/14/Z.

## Appendix

**Supplementary Figure 1:**
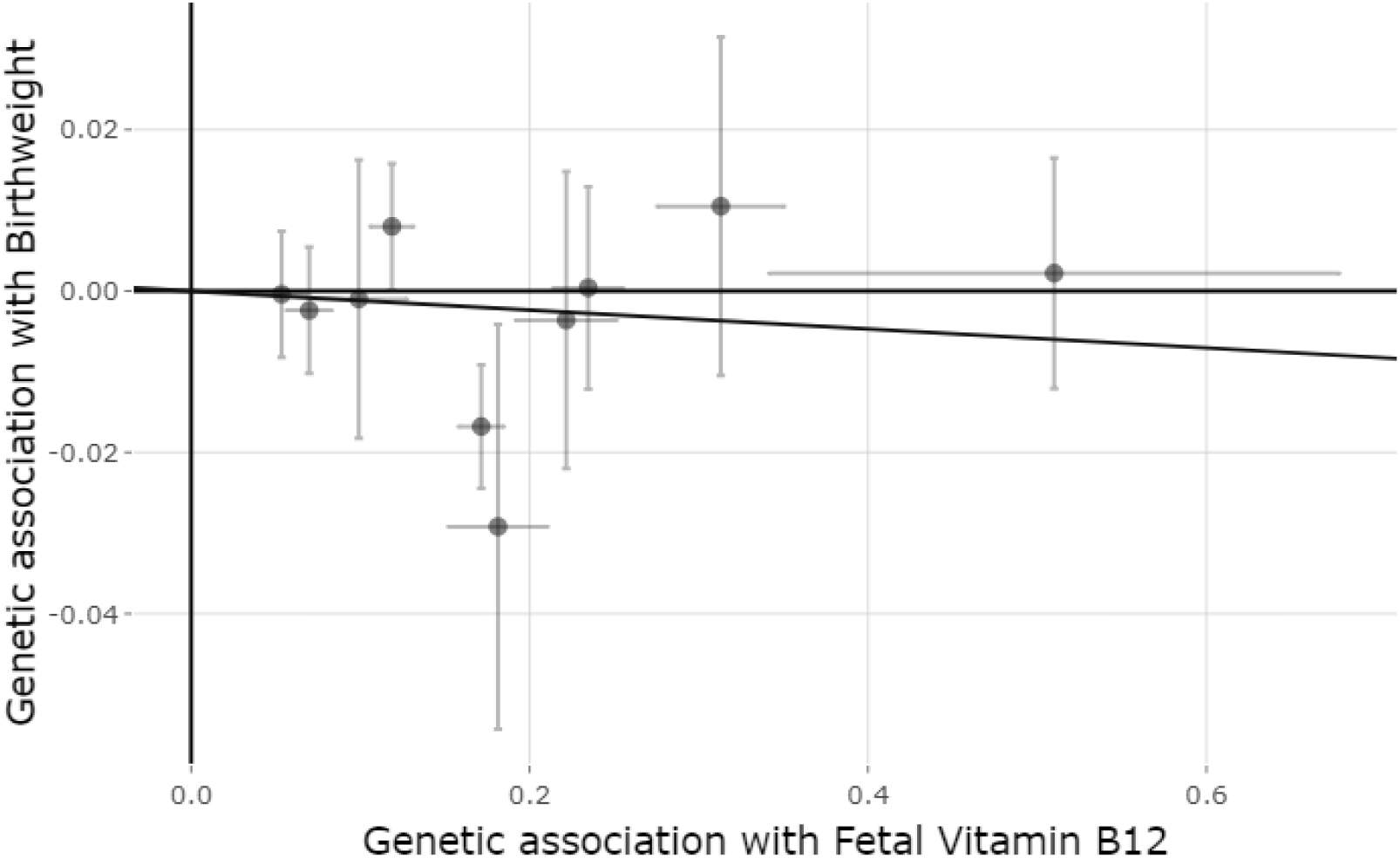
Mendelian randomization estimates of the causal effect of fetal vitamin B12 levels on offspring birthweight. X-axis shows the SNP effect, and standard error, on vitamin B12 levels for each of the ten SNPs and Y-axis shows the SNP effect, and standard error, on offspring birthweight. The regression line for the IVW MR method is shown.

**Supplementary Figure 2:**
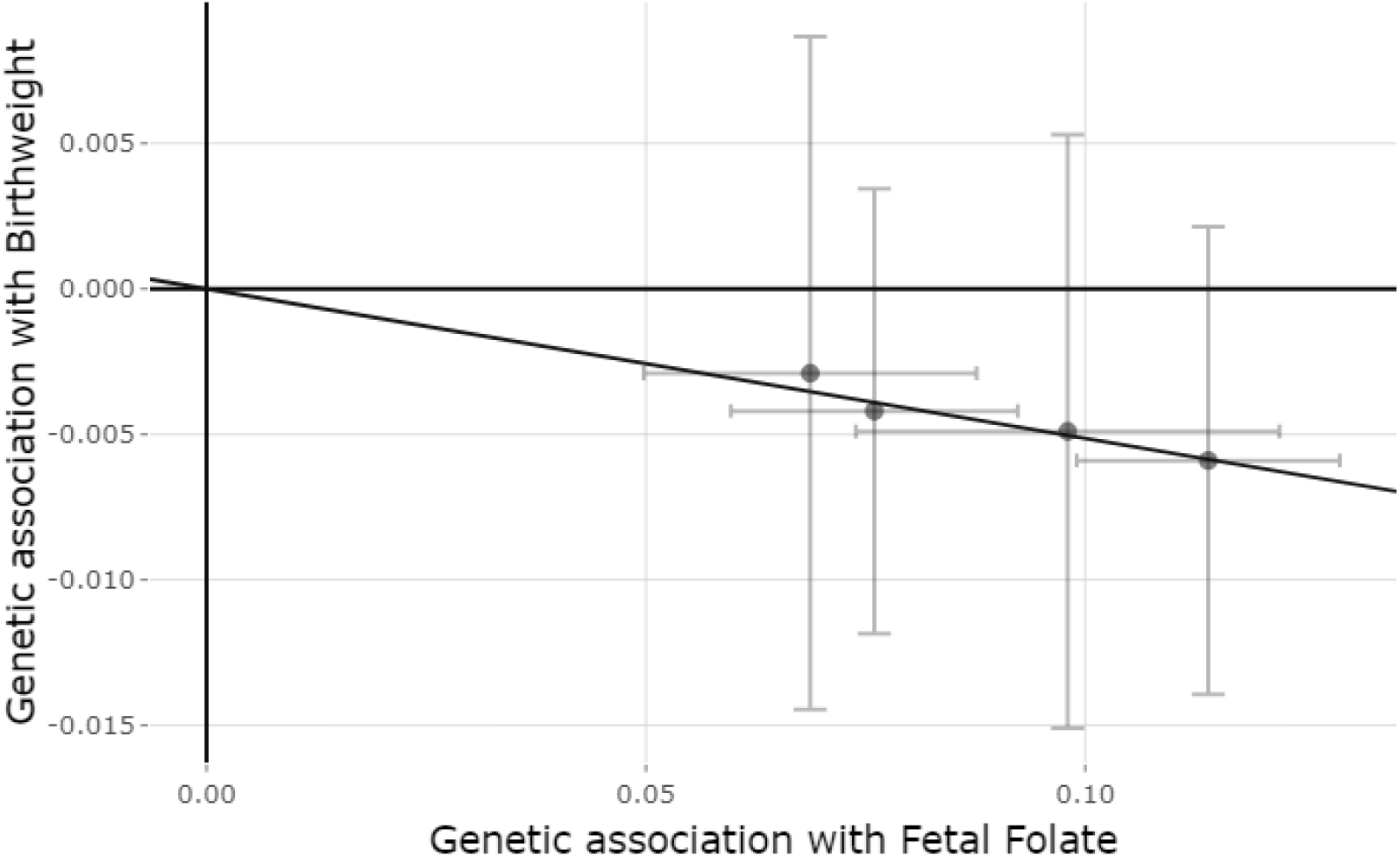
Mendelian randomization estimates of the causal effect of fetal folate levels on offspring birthweight. X-axis shows the SNP effect, and standard error, on folate levels for each of the four SNPs and Y-axis shows the SNP effect, and standard error, on offspring birthweight. The regression line for the IVW MR method is shown.

**Supplementary Figure 3:**
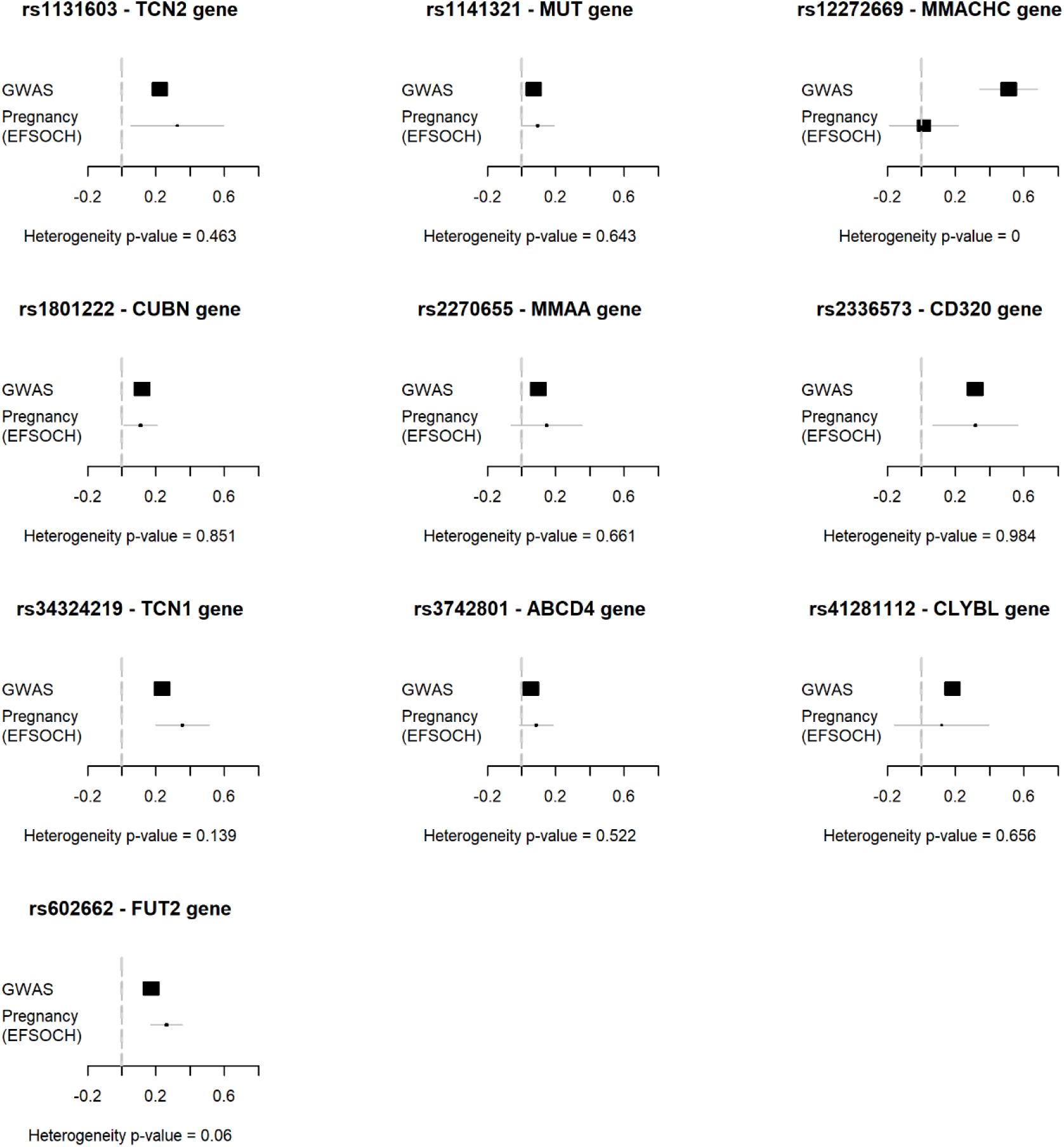
Forest plot comparing the SNP effect sizes on B12 from the exposure GWAS used in the MR analysis with the corresponding SNP effects on B12 measured in pregnant women from the EFSOCH study. All plots show overlapping confidence intervals and no sign of heterogeneity except for in the rs12272669 variant in the *MMACHC* gene (heterogeneity p = 0.0002)*. *We used a proxy for this variant in the EFSOCH study, rs11234541, which was in high LD with rs12272669 but had low imputation quality.

**Supplementary Figure 4:**
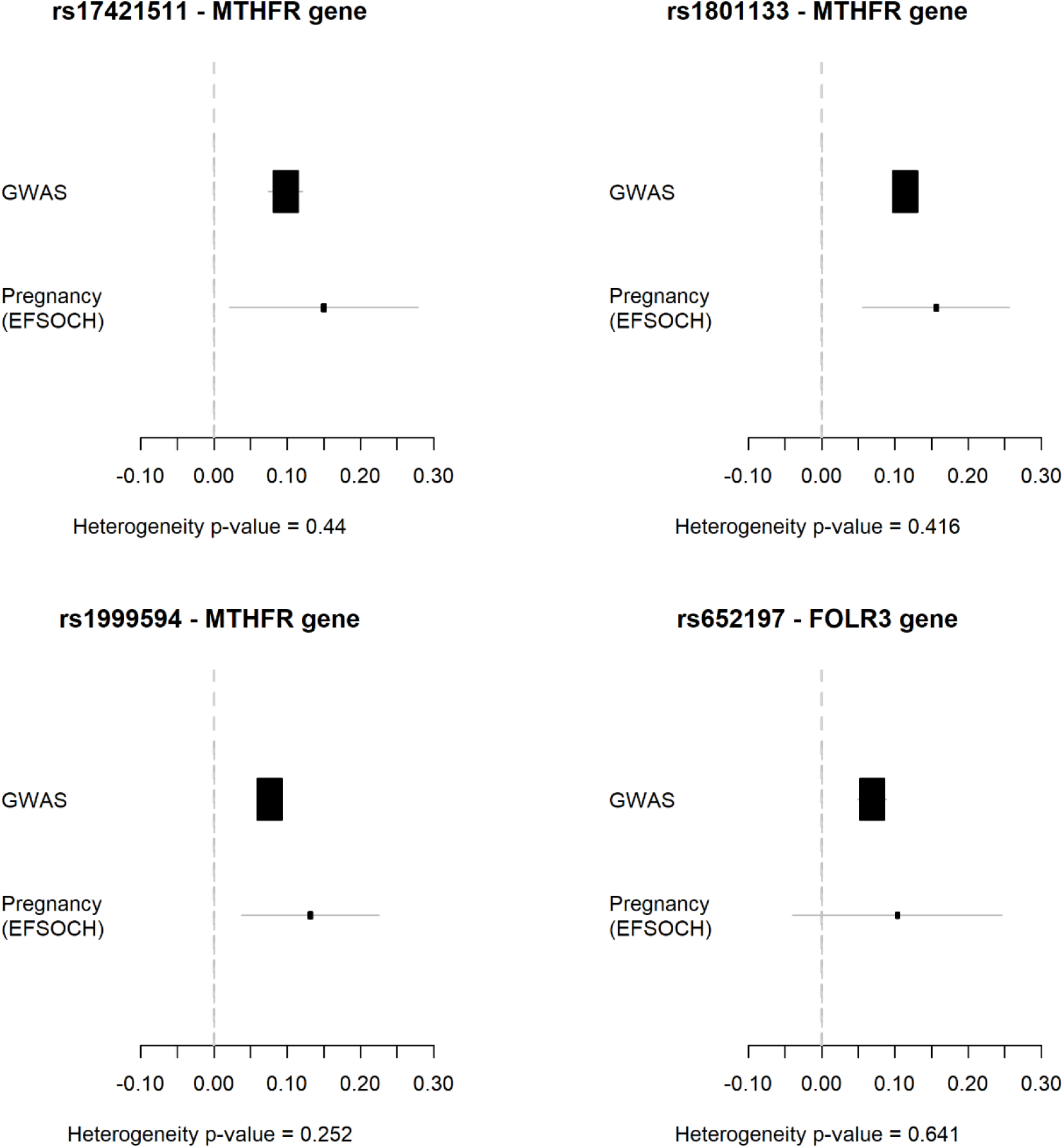
Forest plot comparing the SNP effect sizes on folate from the exposure GWAS used in the MR analysis with the corresponding SNP effects on folate measured in pregnant women from the EFSOCH study. All plots show overlapping confidence intervals and no sign of heterogeneity.

